# casmini-tool: a comprehensive database for efficient and specific guide RNA design using dCasMINI

**DOI:** 10.1101/2023.09.17.558168

**Authors:** Spencer Lopp, Tyler Borrman, Nishant Jha, Melanie Silvis, Ryan Swan, Xiao Yang, Gabriella Alvarez, Courtney Klappenbach, Yanxia Liu, Daniel O. Hart, Lei S. Qi, Timothy P. Daley, Robin W. Yeo

## Abstract

The dCasMINI protein is a hypercompact, nuclease-inactivated CRISPR-Cas system engineered for transcriptional modulation and epigenetic editing [Xu et al., 2021]. The small size of dCasMINI (529 amino acids), less than half the size of comparable Cas9 molecules, makes it ideal for AAV-based therapies which are frequently limited by AAV’s small cargo capacity. Unlike Cas9 or Cas12a, there are no available computational tools for designing dCasMINI guides. To facilitate and accelerate the development of dCasMINI-based applications, we synthesized knowledge regarding dCasMINI guide design and built a website to assist researchers in designing optimal guides for dCasMINI-based experiments for transcriptional inhibition (CRISPRi) and activation (CRISPRa); to ensure that our tool would be useful for therapeutic guide design, in which a guide’s off-target safety profile is of paramount importance, we specifically optimized alignment parameters for high-sensitivity to comprehensively report genome-wide off-targets. To investigate dCasMINI’s full protospacer adjacent motif (PAM) profile, we engineered libraries of PAMs and exhaustively characterized dCasMINI’s ability to activate a locus with different PAMs. We also experimentally investigated the importance of each nucleotide position on the guide RNA’s ability to activate its target, and characterized a 6bp high-fidelity seed region at the 5’ end of the protospacer sequence which we identified to be intolerant to mismatches and deletions, and thus critical for true binding events. Taken together, our tool offers CRISPRi/a guide design for every protein-coding gene in the human genome along with comprehensive off-target prediction, incorporating the most up-to-date information about dCasMINI’s full PAM and protospacer design rules. The tool is freely available to use at www.casmini-tool.com.

---

The CRISPR-Cas revolution is upon us. Advances in CRISPR-Cas9 based therapeutics have resulted in transformational therapies for *β*-thalassemia [Frangoul et al., 2020], sickle cell disease [De Dreuzy et al., 2019], B-cell lymphoma [McGuirk et al., 2022], non-Hodgkin lymphoma [O’Brien et al., 2022], and hereditary transthyretin amyloidosis [Gillmore et al., 2021], among others. However, the large size of the Cas9 molecule presents challenges for therapeutic delivery. For example, its large size (in the range of 3-4kb) limits its use with adeno-associated virus (AAV) delivery, which has a packaging size below 4.7kb [Wu et al., 2010]. As a consequence, the vast majority of current CRISPR-Cas9 therapies are restricted to ex-vivo or lipid nanoparticle-based delivery modalities, severely limiting their general application. The dCasMINI molecule [Xu et al., 2021] is ideal for AAV-based delivery, as its small size of 529 amino acids (1587bp) allows researchers to package the DNA sequence for dCasMINI fused to a functional peptide or enzymatic domain capable of gene regulation (which we term “modulator”), along with the promoter sequence and guide RNA sequence, into a standard AAV vector (maximum cargo size of 4.7kb). Furthermore, traditional Cas9-based therapeutics are limited to the treatment of diseases that are ameliorated by gene knockouts and are thus unsuitable for a whole host of genetic diseases such as those caused by haploinsufficiency. On the other hand, dCasMINI has an inactivated nuclease domain and is thus capable of upregulating or downregulating a genetic locus when tethered to the appropriate modulator peptides. Beyond its smaller size and modulatory versatility, recent research has additionally suggested that dCasMINI has a lower incidence of off-targets than Cas9 or Cas12a [Xin et al., 2022], making it an attractive Cas molecule candidate for therapeutic applications.

To assist researchers in the application of dCasMINI-based tools, we have developed a web-based database to rapidly search for optimal dCasMINI spacer sequences for CRISPR interference (CRISPRi) and CRISPR activation (CRISPRa) at any desired genetic locus in the human genome (Figure 1). Virtually all Cas proteins require a PAM sequence flanking the DNA target to allow for binding and subsequent activity. Depending on the specific Cas protein, these PAMs vary in sequence, length, orientation, and target distance [Collias and Beisel, 2021]. Using a PAM library screen we extensively characterized the PAM recognition profile for dCasMINI to increase guide coverage and off-target sensitivity. To improve upon pre-existing tools such as CHOPCHOP [Labun et al., 2019] or Casilico [Asadbeigi et al., 2022], we performed extensive computational mapping of each potential spacer sequence with high-sensitivity parameters to extend the search for off-targets, and provided the full off-target information for each guide so that researchers can determine what type and how many off-targets they can tolerate and/or test. To further characterize computationally predicted off-targets, we experimentally determined the guide seed region (high-fidelity seed region at 5’ of spacer that is intolerant to mismatches and deletions) for dCasMINI and annotate potential off-targets with information about seed region mismatches. This will allow researchers to increase the potential search space of guides by including guides with computationally predicted off-targets that have mismatches in the seed region (and thus unlikely to result in a true binding event). We expect that our casmini-tool and website portal will greatly simplify and accelerate guide design workflow for researchers carrying out experiments using the dCasMINI platform.

**Figure 1:**
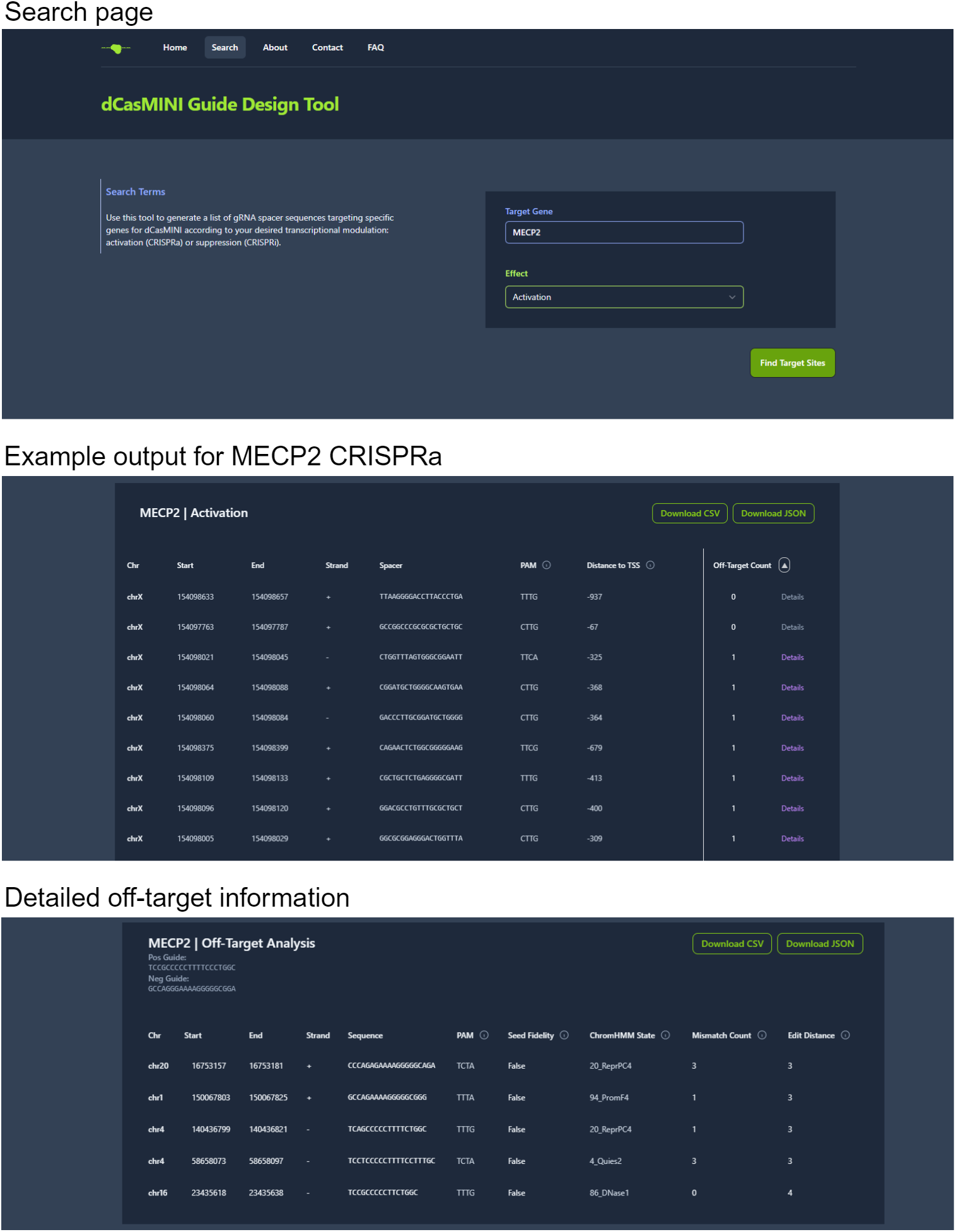
Example workflow for designing guides targeting the Rett syndrome associated gene MECP2 for CRISPRa.

## Methods

### dCasMINI PAM profile characterization

To determine which PAMs allow for dCasMINI binding, we developed a pooled PAM screening assay using a synthetic reporter, flow cytometry, and NGS to evaluate all possible nucleotide arrangements of the PAM. Briefly, a pooled library of synthetic PAM reporter lentivirus was designed such that a variable PAM region of 6 random nucleotides (N6) was adjacent to a fixed synthetic protospacer binding sequence, and placed directly upstream of a weak synthetic promoter driving expression of GFP. The reporter also encodes for the constitutive expression of BFP. The lentiviral reporter library was stably integrated into HEK293T cells via transduction such that the majority of cells receive a single reporter copy (therefore a single PAM sequence), and followed by transduction of the GEMS components: constitutive expression of dCasMINI-EpiMod (strong activator) [Carosso et al., 2023] and either the targeting sgRNA or a non-targeting sgRNA. We used FACS to separate GFP+ and GFP-cells, and generated NGS libraries from each population using PCR to amplify the PAM region of the reporter. We used DESeq2 to assess the enrichment of each N6 PAM in the GFP+ population compared to the GFP-population, and defined significant hits using the non-targeting sgRNA library controls (Log2FC>1.5, padj<0.01). To define the N4 PAM hits, we looked for consistency within the N6 hits (>7/16 N6 hits).

### Computational identification of on-target gRNA sequences

To identify on-target guide sequences for CRISPRi/a genome-wide, we first extracted the primary transcription start site (TSS) for each gene using the hg38 FANTOM5 database, a repository of genome-wide TSSs defined from human CAGE-seq data [FANTOM, 2014]. Filtering out mitochondrial genes resulted in a dataset of 22,499 genes. Upon inspection, several known protein-coding genes were missing from the FANTOM5 database. To correct for this, we intersected the FANTOM5 genes with the GENCODE v45 set of protein coding genes to discover an additional 2,666 genes annotated by GENCODE but missing from FANTOM5 [Frankish et al., 2021]. The TSS for these supplemental GENCODE genes were derived from the gene start locations of the GENCODE GTF. In total a set of 25,165 genes with TSS coordinates were included in the database. Based on in-house experiments [data not shown], we defined the optimal targeting region for each regulatory modality as follows:

- CRISPRi: -200bp:+1000bp from the TSS;
- CRISPRa: -1000bp:+200bp from the TSS.

Nucleotide sequences for each of these targeting regions genome-wide were extracted using the pybedtools command getfasta [Dale et al., 2011]. We then identified all dCasMINI PAMs (TTTA, TTTG, TCTA, CTTG, TTCA, TTCG) in these sequences and extracted the 20 nucleotides directly downstream of each PAM to generate the list of on-target spacer sequences.

### Computational mapping of putative off-targets

We mapped each 20bp on-target spacer sequence to the hg38 genome with Bowtie2 [Langmead and Salzberg, 2012], modifiying Bowtie2’s –very-sensitive preset option to further increase sensitivity in off-target predictions with a maximum of twenty thousand alternative mapping locations, using the following command:

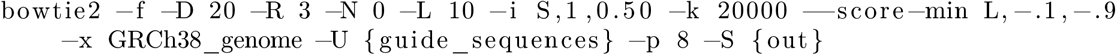

For each putative off-target site, we queried 4bp upstream and discarded any alignments that did not contain an appropriate PAM in the upstream region. Given our experimental data on the importance of the 6bp high-fidelity seed region (see below), we further verified whether any mismatches or deletions were present in the first 6bp of the 20bp spacer alignment and annotated each off-target with this information.

### Experimental determination of the dCasMINI seed region

To identify the high-fidelity seed region of dCasMINI spacer sequences, we synthesized a panel of gRNA-encoding gene fragments where each individual nucleotide in the spacer sequence targeting the CD2 locus was mutated to all three other nucleotides (“single-mismatch”) or deleted (“single-deletion”). Each gRNA variant was cloned into the gRNA plasmid backbone downstream of mU6 promoter. HEK293T cells were co-transfected with individual gRNA variant plasmids and dCasMINI-EpiMod (strong activator) [Carosso et al., 2023] as triplicates in a 96-well plate format. Three days post-transfection, CD2 target gene activation was quantified by cell surface antibody staining of live cells using APC anti-human CD2 antibody (Biolegend, 309224) followed by flow cytometry on the cell population transfected with both gRNA and dCasMINI plasmids (performed on Cytoflex and analyzed using Flowjo software). The level of gene activation by each gRNA variant was compared and normalized to the activity of the wild-type gRNA as a control. The seed region in the spacer with minimal tolerance on single mismatch/deletion was defined with a cutoff of normalized activity below 0.2.

### ChromHMM annotation of off-targets

We employed a full stack ChromHMM model providing universal chromatin state annotations of the human genome to annotate off-targets with their predicted epigenomic state [Vu and Ernst, 2022]. ChrommHMM state annotations were intersected with off-target coordinates using bedtools intersect [Quinlan and Hall, 2010].

## Results

### Case study: generating CRISPRa guides against MECP2

As a case study, we queried our website tool for CRISPRa guides against MECP2, a gene frequently mutated in Rett syndrome, a severe and progressive X-linked neurodevelopmental disorder (Figure 1) [Lamonica et al., 2017]. Querying “Target Gene” = MECP2 and “Effect” = Activation on the website portal generates a list of on-target guide sequences situated -1000bp:+200bp around the TSS (since that is the appropriate targeting window for transcriptional activation). On-target guide sequences are displayed in a custom genome viewer illustrating their positioning within the chromosome and their proximity to their associated gene’s TSS. The custom genome viewer can be toggled to IGV allowing for a more in-depth visualization of guide sequences [Robinson et al., 2023] (Figure 2). On-target spacer sequences are annotated with chromosomal position, strand, upstream PAM, distance to *the TSS*, and off-target count. These guides are ranked in ascending order based on number of off-targets to prioritize the more therapeutically relevant guide sequences with few predicted off-targets. When interested in a specific spacer sequence, the user can further query information about predicted off-target binding sites by following the “Details” link beside the “Off-Target Count” column (Figure 1).

**Figure 2:**
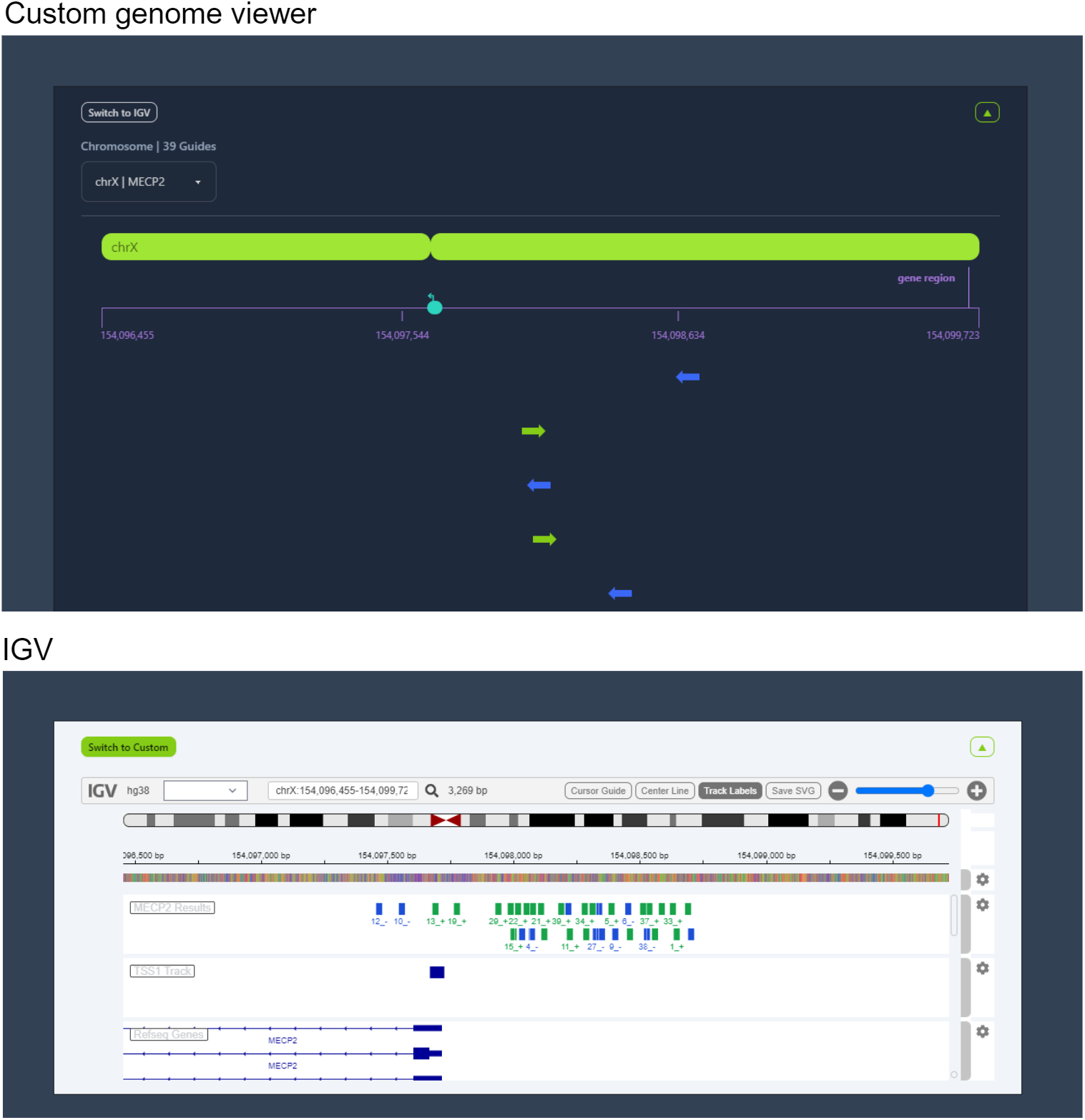
Custom genome viewer and IGV visualization of CRISPRa guides targeting MECP2

### dCasMINI PAM profile characterization

Relative to Un1Cas12f1 from which dCasMINI was derived, dCasMINI contains mutations in the PAM recognition region allowing for possible recognition of alternative PAM sites in addition to the known canonical TTTR PAMs (TTTA and TTTG) of Un1Cas12f1. To investigate dCasMINI’s full PAM profile, we engineered libraries of all possible nucleotide arrangements of a 6 bp motif (4^6^ = 4096 random PAMs). This N6 library was used to screen for PAM sequences which allow for dCasMINI binding in a GFP+ versus GFP-reporter assay. Conservation of the 4 bps of the N6 PAM nearest the spacer sequence led us to identify PAM sequences of 4 bp in length allowing for dCasMINI recognition. Bioinformatic analysis revealed that in addition to the two canonical TTTR PAMs, dCasMINI was found to recognize several other additional “relaxed” PAMs (TCTA, CTTG, TTCA, TTCG), less potent than TTTR but sufficiently strong for target regulation (Figure 3). We incorporated canonical and relaxed PAMs into our guide design algorithm to expand the set of candidate on-target guides and increase the sensitivity of off-target prediction.

**Figure 3:**
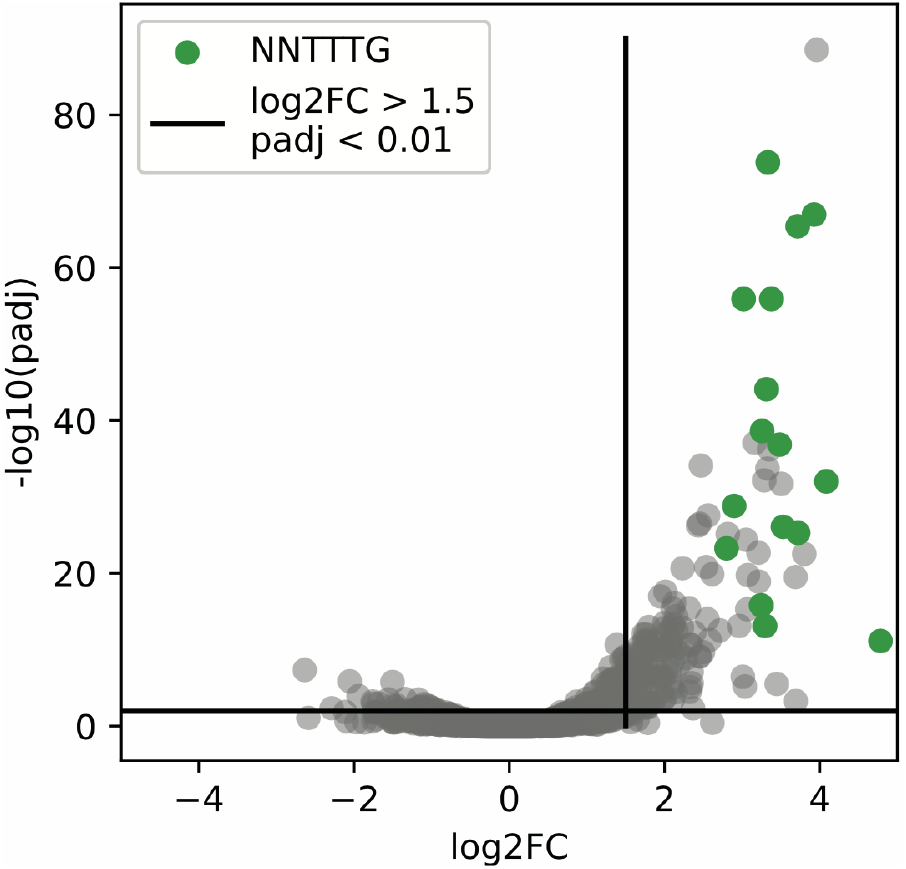
Volcano plot illustrating results of N6 PAM screening. Top right quadrant represents statistically significant N6 PAMs enriched in GFP+ libraries.

### A comprehensive database for computational identification of on-target gRNA sequences

Our tool provides on-target dCasMINI guide sequences for 25,165 human genes suitable for CRISPRi and/or CRISPRa (based on position relative to the primary TSS). In total, we identified 1,172,461 spacer sequences suitable for CRISPRi and 1,267,870 spacer sequences for CRISPRa (347,368 spacer sequences were situated +/-200bp around the TSS and thus appropriate for both CRISPRi and CRISPRa). For CRISPRi, genes have an average of 47 on-target spacer sequences suitable for transcriptional suppression (Figure 4). For CRISPRa, genes have an average of 50 on-target spacer sequences suitable for transcriptional activation (Figure 5).

**Figure 4:**
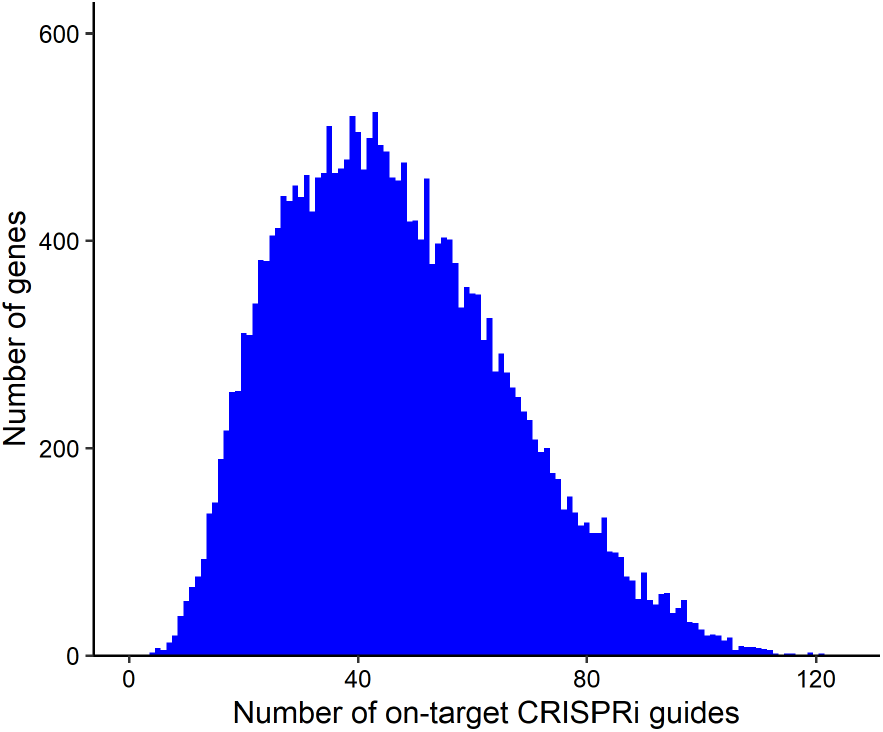
Histogram displaying number of on-target CRISPRi guides per gene genome-wide.

**Figure 5:**
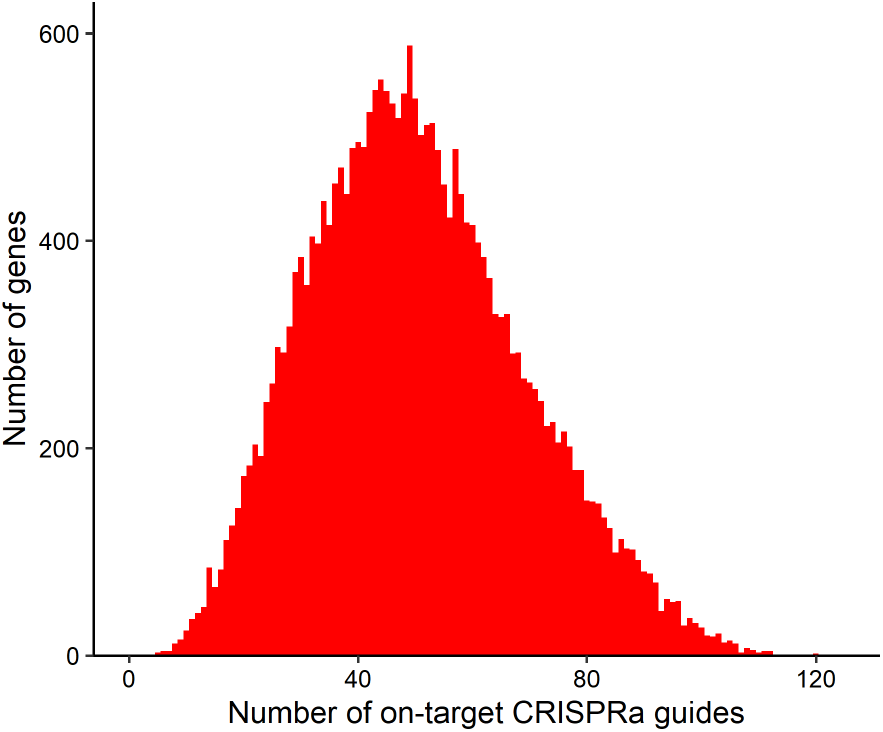
Histogram displaying number of on-target CRISPRa guides per gene genome-wide.

### dCasMINI seed region experiment

To computationally predict off-targets, we first investigated the tolerance of dCasMINI on mismatched target sites and characterized the high-fidelity seed region on its spacer sequence [Slaymaker et al., 2016]. We systematically mutated the CD2 guide spacer sequence to introduce single-base mismatches and single-base deletions at different positions, and then measured the tolerance of dCasMINI-EpiMod against these spacer variants in the context of CRISPRa. We identified a conserved high-fidelity seed region in the +1 to +6 range where any single mismatches or deletions functionally abolish CRISPRa activity (Figure 6).

**Figure 6:**
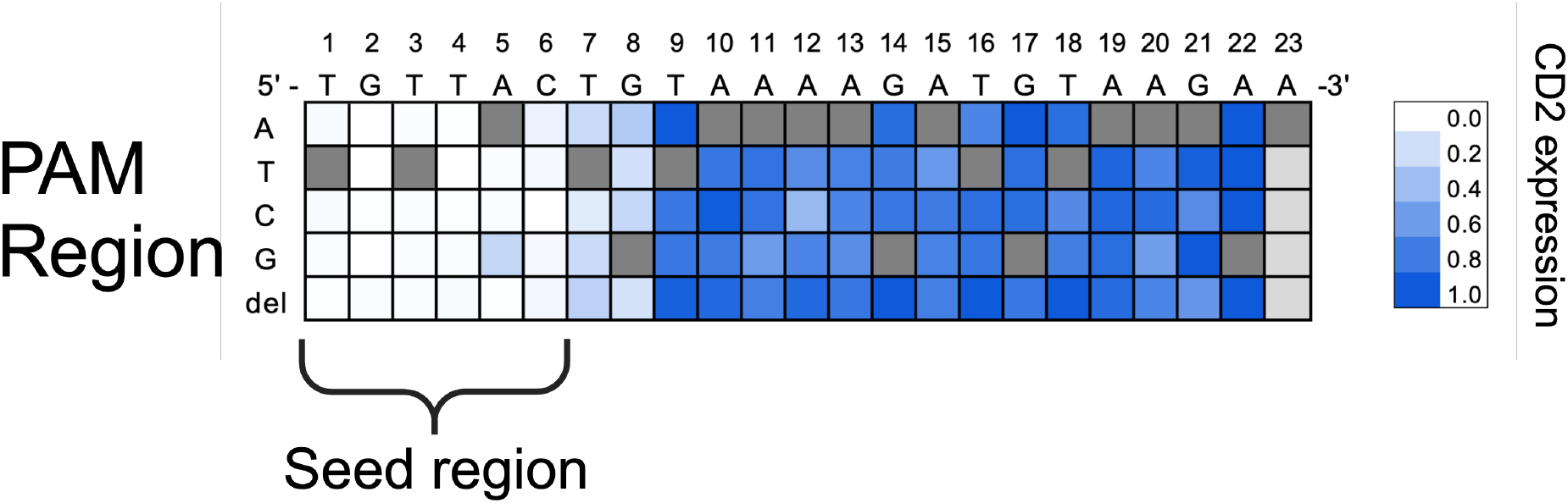
Heatmap illustrating single mismatch/deletion tolerance of dCasMINI spacer sequences targeting the endogenous CD2 locus for transcriptional activation, 3 day post transfection. The heatmap was generated by normalizing CD2 expression with a given single mismatch/deletion spacer to the maximum expression level of CD2 with the WT gRNA sequence. Lower expression indicates that the mismatch disrupts binding.

### Computational mapping of putative off-targets

To provide comprehensive off-target information for CRISPRi/a guides, we used Bowtie2 to sensitively map each on-target spacer sequence genome-wide and annotated potential off-targets with information about whether mismatches or deletions were present in the high-fidelity 6bp seed region of the spacer. We anticipate this will allow researchers to prioritize testing guides with minimal predicted off-target effects, which is critical for developing safe and efficacious therapeutic products. In total, we mapped 1,382,617,925 potential off-target sites associated with CRISPRi/a spacer sequences.

Ultimately we predict users will be most interested in selecting guides with a minimal number of potential off-target binding events. Analyzing the off-target database reveals that genes have an average of 22 on-target guides with ≤ 5 potential off-target sites for CRISPRi (Figure 7) and 21 on-target guides with ≤ 5 potential off-target sites for CRISPRa (Figure 8). However, many of these potential off-target sites (approximately 26%) contain one or more mismatches in the spacer high-fidelity seed region which are extremely unlikely to result in true off-target binding events (Figure 6). Incorporating these data on seed region fidelity (removing off-targets with mismatches in the seed region) we increase the average on-target guides per gene to 33 and 34 for CRISPRi and CRISPRa, respectively (Figure 7 and Figure 8). An off-target’s capacity to cause harm is linked to its genomic context. For example, an off-target located in a silent region of the genome devoid of regulatory sites is likely to be less hazardous than an off-target located at a gene promoter or enhancer region. To incorporate this information, we annotated each of the putative off-target sites with predicted chromatin state annotations from a universal full stack ChromHMM model [Vu and Ernst, 2022]. We expect these chromatin state annotations to further guide users in determining whether predicted off-targets will be detrimental to their specific studies.

**Figure 7:**
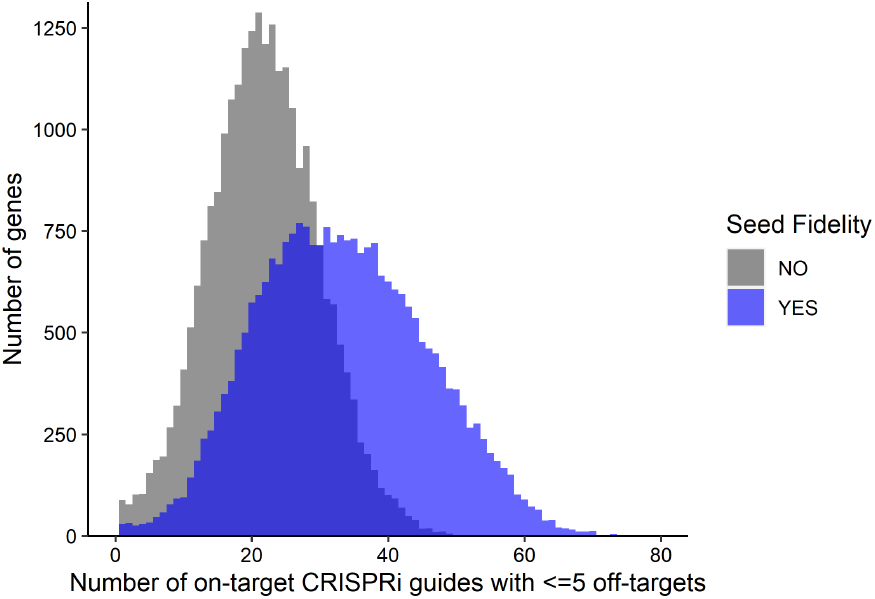
Histograms displaying number of on-target CRISPRi guides with 5 potential off-target sites per gene genome-wide. Seed Fidelity YES category restricts off-targets to those with seed fidelity (no mismatches in seed region). Seed Fidelity NO category sets no restriction on seed fidelity (off targets may or may not contain mismatches in seed region).

**Figure 8:**
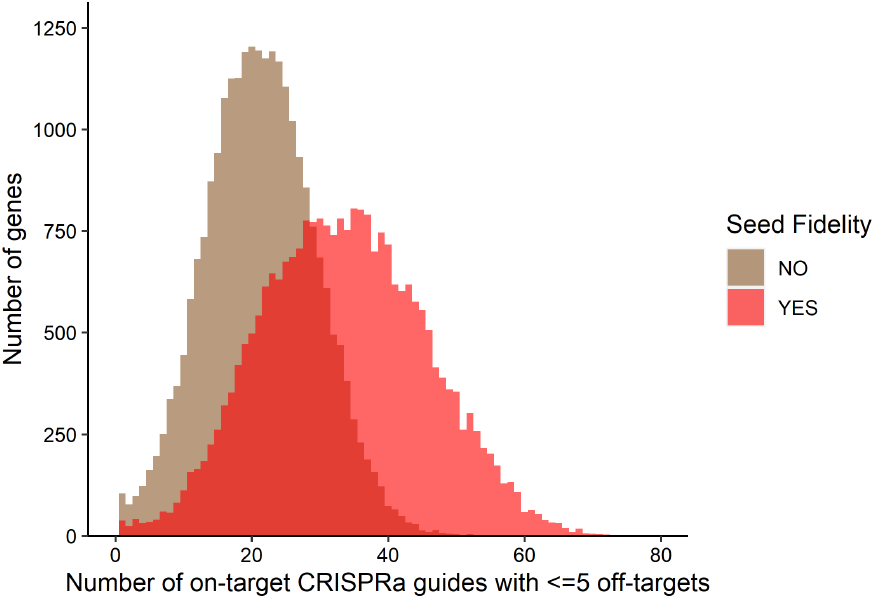
Histogram displaying number of on-target CRISPRa guides with 5 potential off-target sites per gene genome-wide. Seed Fidelity YES category restricts off-targets to those with seed fidelity (no mismatches in seed region). Seed Fidelity NO category sets no restriction on seed fidelity (off targets may or may not contain mismatches in seed region).

## Discussion

The simplicity and versatility of the CRISPR-Cas9 platform for genetic editing has increasingly resulted in its use as a therapy for the treatment of genetic diseases [Wang et al., 2016]. While extremely powerful, the traditional CRISPR-Cas9 platform suffers from a number of key drawbacks: notably, its large size prevents it from being delivered in a single AAV and its nuclease ability renders it unsuitable for the treatment of a host of genetic disorders in which healthy genes are inactivated or more subtle downregulation is required. Since the discovery of Cas9 [Jinek et al., 2012], there has been an explosion of research into the diversification and optimization of other Cas molecules [Chavez et al., 2023]. Recently, a hyper-compact, nuclease-inactivated Cas molecule, termed dCasMINI, was engineered to be small enough for AAV delivery, while maintaining high on-target efficiency and specificity, making it an ideal Cas molecule candidate for the development of AAV-based therapies [Xu et al., 2021]. When fused to a transcriptional activator, dCasMINI can drive thousand-fold increases in target gene activation [Xu et al., 2021]. Such functionality is critical for targeting diseases caused by haploinsufficiency or loss of function mutations which traditional Cas9 nuclease approaches cannot address. For example, Rett syndrome is caused by a loss of function mutation in the gene MECP2 [Amir et al., 1999]. Considering dCasMINI’s capability for strong target gene activation, dCasMINI mediated CRISPRa of the healthy MECP2 allele may be a potential therapeutic strategy to treat Rett syndrome patients.

As different Cas molecules are governed by different guide design principles, guide sequence design is a frequent bottleneck in the dCasMINI workflow for CRISPRi/a. Here we present a tool to easily design gRNAs against any human loci of interest for CRISPRi/a experiments using dCasMINI. We expect the full PAM profile characterization and mismatch data in the high-fidelity seed region to further enhance gRNA selections compared with current tools supporting type V CRISPR-Cas systems such as Casilico [Asadbeigi et al., 2022] and DeepCpf1 [Kim et al., 2018]. In comparison to off-target algorithms employed by guide design tools such as CHOPCHOP [Labun et al., 2019] and CRISPOR [Concordet et al., 2018] which do not support gapped alignments, casmini-tool more exhaustively characterizes the off-target space by reporting off-targets containing gaps and/or mismatches (known to be tolerated to some extent by dCasMINI and other Cas molecules [Lin et al., 2014]). Considering detrimental off-target events are likely linked to genomic/epigenomic context, the presented chromatin state annotations also provide further insight into predicted gRNA safety.

In summary, in this study we present a novel tool for dCasMINI gRNA design, comprehensive characterization of potential off-targets, and detailed visualizations of both on-target and off-target results. We anticipate that this web-based tool will be of value to the CRISPR-Cas research community and will allow researchers to more easily and quickly design CRISPRi/a experiments with dCasMINI.

## Acknowledgements

The authors would like to thank employees of Epicrispr Biotechnologies for helpful conversation and feedback during the course of this project.

## Authors’ Contributions

S.L., T.B., and R.W.Y. developed the backend database of on- and off-target guides. S.L. and N.J. built the frontend website. M.S. and R.S. developed and analyzed the PAM characterization experiments. X.Y., G.A., and C.K. developed and analyzed the seed region experiments. T.P.D. and R.W.Y. supervised and directed this project. R.W.Y., T.B., and T.P.D. wrote the manuscript with input from all authors.

